# A scalable proteogenomic framework for dissecting phospho-signaling pathways in primary immune cells

**DOI:** 10.1101/2025.10.08.681012

**Authors:** Christian M. Beusch, Carolyn Morningstar, Marc Semaan, Christopher M. Monaco, Sarah Welbourn, Hailey Swaldi, Jae-Kyun Ko, Devin Kenney, Alexander Sousa, Holt Sakai, David Liu, Hisashi Akiyama, Mohsan Saeed, David E. Gordon

## Abstract

Signaling networks modulated by post-translational modifications orchestrate cellular responses to external cues. Traditional approaches to study these pathways lack the throughput to systematically capture the causal architecture of these signaling pathways at scale. Here, we present an integrated proteogenomic framework that combines saturating genetic perturbations with high-throughput proteomics to systematically map cytokine-induced signaling in primary human T cells. Supporting this framework is *simplePhos*, a streamlined, low-input phosphoproteomics workflow that enables scalable, time-resolved analysis without the requirement for specialized equipment or robotics. We extensively validate the *simplePhos* pipeline by applying inflammatory stimuli, including type I and II interferons, lipopolysaccharide, and Sendai virus to primary T cells and myeloid cells, establishing foundational datasets in these treatment contexts. Ultimately, using type I interferon signaling in genetically modified T cells as a model, we demonstrate that combined application of genetic alterations and proteomic analyses can map key signaling nodes in primary immune cells. This represents a powerful strategy to mechanistically interrogate phospho-signaling networks in human immune cells, with broad applications in translational immunology and therapeutic development.

## Introduction

Cellular signaling involves dynamic changes in covalent and non-covalent biochemical interactions, with proteins acting as the effectors of these regulatory processes. Specifically, changes in protein post-translational modifications (PTMs) are central to signal transduction, ultimately influencing protein expression, cellular behavior, and physiological responses^1, 2^. Mass spectrometry-based proteomics has become a powerful tool for studying these signaling events, providing system-wide resolution of PTM dynamics and protein levels in response to external stimuli or disease-related disturbances. However, in the past, proteomics-driven studies of signaling networks have focused primarily on observational time-course analyses, which capture temporal changes in protein modifications and abundance following stimulation^3, 4, 5^. While such studies are instrumental for charting stimulus-associated responses, they are inherently limited in their ability to establish causal relationships within signaling networks. Small-molecule agonists and inhibitors have traditionally been used to identify key nodes in signaling pathways; however, their utility is constrained by limited availability and frequent off-target effects, particularly among kinase inhibitors^6, 7, 8^. These limitations underscore the need for complementary strategies with greater specificity and scalability. The integration of genetic perturbations, particularly loss-of-function approaches, into proteomics analyses provides a compelling avenue for causal inference, enabling systematic dissection of signaling hierarchies and regulatory nodes. Applying this strategy, we developed a proteogenomic framework that combines high-throughput global abundance and phosphoproteomics with targeted genetic perturbations, applying both saturating knockouts and base editing, to systematically map causal elements of signaling pathways in primary human immune cells^9, 10^.

In this study, we focused on type I interferon (IFN-I) signaling as a model system due to its well-characterized components and proteomic phenotypes, such as JAK/STAT-mediated phosphorylation and interferon-stimulated gene (ISG) induction^11, 12^. IFN-I signaling intersects with key pathways involved in inflammation, viral defense, cancer, autoimmunity, and interferonopathies, making it a particularly relevant axis for detailed signaling dissection^13, 14, 15^. Given that innate immune pathways are often dysregulated in cancer and other diseases, transformed cell lines may not faithfully recapitulate physiological signaling^16, 17, 18^. To overcome this, we employed primary human CD4 T cells, derived from peripheral blood, as a physiologically relevant model system. CD4 T cells are not only central regulators of adaptive immunity but are also implicated in interferon-driven pathology across diverse inflammatory and autoimmune conditions^19^. Among PTMs, phosphorylation is key to a multitude of immune regulatory pathways, including interferon signaling. It is rapid, reversible, and capable of inducing substantial conformational changes due to its strong negative charge; therefore, phosphorylation is well-suited for both information transfer and modulation of protein functionality. The availability of known phosphorylation targets downstream of IFN-I signaling, such as STAT1 Y701, STAT3 Y705, and STAT4 Y693, provides a robust reference point for benchmarking perturbation effects and studying regulatory layers^20, 21, 22, 23, 24, 25, 26^. More broadly, by integrating precise genetic perturbations with unbiased proteomics, our approach enables systematic mapping of signaling circuits in primary immune cells, with applications in immunotherapy development, vaccine optimization, and the mechanistic study of immune dysregulation.

## Results

### Mapping the global proteomic ISG landscape in primary human CD4 T cells

Much of our current knowledge of interferon signaling derives from transcriptomic profiling or studies in transformed cell lines, experimental systems that may incompletely capture the proteomic responses of primary human immune cells^27, 28, 29, 30, 31, 32, 33, 34^. Therefore, we employed high-resolution mass spectrometry to systematically profile IFN-I driven protein expression in primary CD4 T cells purified from healthy donors (Figure 1A). CD4 T cells from two healthy donors were stimulated with two concentrations of interferon-β (100 U/ml and 1000 U/ml) for up to 24 hours. Quantitative global proteomic profiling captured up to 8,000 proteins per sample with high reproducibility, enabling comprehensive interrogation of IFN-β–driven responses ( Supplementary Figure 1A). Indeed, by 24 hours post stimulation, we observed robust and reproducible induction of canonical ISGs, including DDX58/RIG-I, MX1, ISG15, and multiple OAS family members across both IFN-β concentrations (Figure 1B)^34^. Notably, the majority of protein-level responses were largely independent of IFN-β dose, with most differential changes already apparent at the lower concentration, suggesting that near-maximal signaling can be achieved with 100 U/ml IFN-β (Figure 1C). Temporal profiling revealed early induction of ISGs by 4 hours post-stimulation, with minimal changes at earlier timepoints (Figure 1D-E, Supplementary Figure 1B). Gene Ontology (GO) enrichment analysis revealed activation of type I interferon signaling pathways, along with additional pathways functionally linked to interferon responses, as early as 4 h post-stimulation. This early enrichment underscores the rapid engagement of antiviral and immune-regulatory programs (Figure 1F, Supplementary Figure 1C). Only a subset of proteins annotated to the “type I interferon signaling pathway” (GO term 0060337) were significantly upregulated in primary CD4 T cells after 24 hours ( Supplementary Figure 1D), likely reflecting our treatment duration, the use of primary CD4 T cells, and other contextual differences from previous interferon studies^28^.

**Figure 1:**
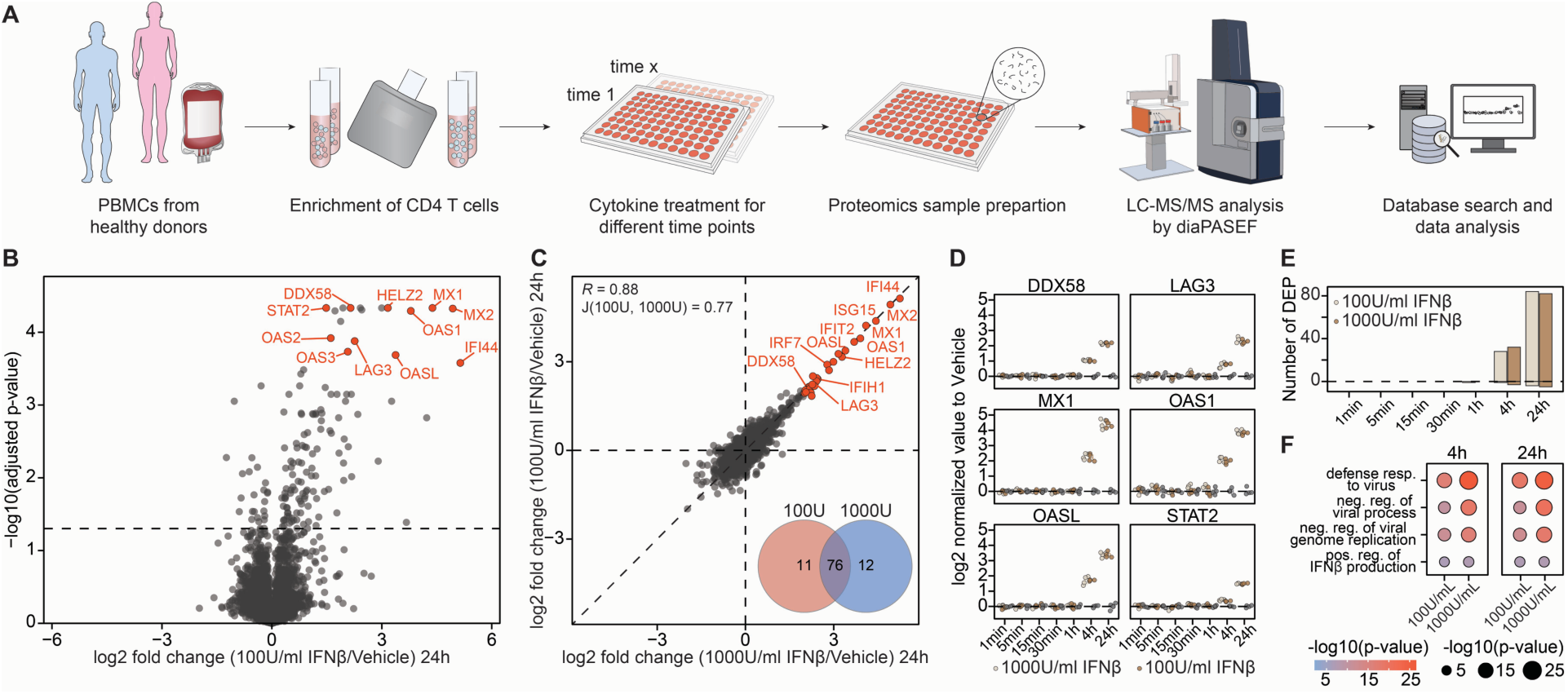
A proteomic atlas of the interferon response in primary human CD4 T cells. (**A**) Schematic overview of the experimental design to profile cytokine-stimulated gene expression via global proteomics in primary immune cells. (**B**) Volcano plot showing differential protein abundance in CD4 T cells 24 hours after treatment with 100 U/mL interferon-β versus vehicle control. (**C**) Scatter plot comparing fold changes in protein expression between 100 U/mL and 1000 U/mL interferon-β treatments in CD4 T cells. A Venn diagram shows the overlap of significantly regulated proteins; Pearson correlation and the Jaccard index quantify similarity between the treatments. (**D**) Boxplots showing temporal expression changes of selected ISGs from panels B and C. (**E**) Bar chart depicting the number of significantly regulated proteins at each time point for both interferon-β concentrations. **(F)** Gene Ontology (GO) enrichment analysis of significantly upregulated proteins upon interferon-β treatment. n = 4, from two donors.

### *simplePhos*: An accessible phosphoproteomics pipeline for signaling analysis at scale

While transcriptomic approaches have advanced our understanding of immune cell states and cytokine responses, elucidating signaling pathways requires scalable, reproducible systems-level measurement of protein phosphorylation. However, there are hurdles towards applying phosphoproteomic workflows to primary samples, including reliance on specialized instrumentation or high sample requirements, factors that limit throughput and accessibility^4, 35, 36, 37, 38^.

To mitigate these limitations and enable scalable phosphoproteomic analyses across diverse timepoints and interventions, we developed *simplePhos*, a streamlined phosphoproteomics workflow built entirely from standard reagents and equipment, enabling high-throughput analysis of low-input primary samples. For phosphopeptide enrichment, we selected Zr-IMAC HP beads, having benchmarked them against Ti-IMAC HP beads and observed comparable yields of localized phosphosites, quantitative accuracy, and phospho-enrichment purity, but superior shelf stability ( Supplementary Figure 2A-C). To evaluate analytical depth, we titrated input amounts from 40 µg down to 2.5 µg of peptides, maintaining a constant bead volume across conditions. We quantified almost 15,000 localized phosphopeptides with reproducibility in higher input samples, and even at the lowest input amount (2.5 µg), we still identified almost 10,000 localized Class I phosphopeptides (localization probability > 0.75), with >90% enrichment purity and minimal loss in quantitative accuracy across the dilution series (Figure 2A, Supplementary Figure 2D). While the number of total phosphopeptides exceeded localized sites ∼4-fold, we attempted to narrow this gap by reducing acquisition windows, a strategy known to enhance phosphosite localization ( Supplementary Figure 2E). To this end, we implemented the recently developed synchroPASEF workflow, though it did not match the depth of localization achieved by our standard DIA workflow^39^ ( Supplementary Figure 2F). We also noticed that the Whisper Zoom liquid chromatography method resulted in an improved number of quantified phosphorylated peptides compared to the legacy Whisper method ( Supplementary Figure 2G-H). This trend is especially noticeable for lower input amounts, and motivated us to use the Whisper Zoom gradient throughout our study (see also Materials and Methods).

**Figure 2:**
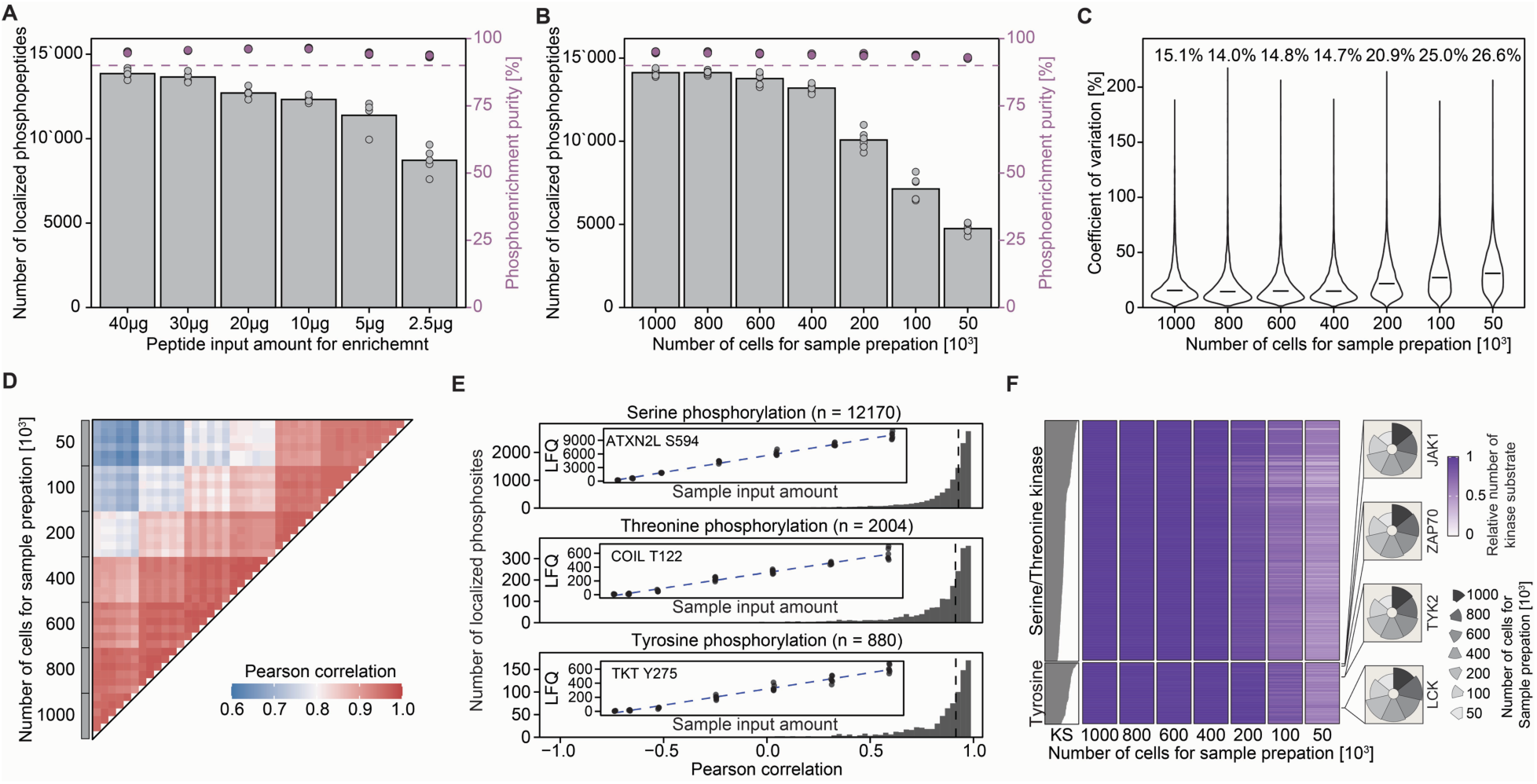
*SimplePhos* enables robust high-throughput phosphopeptide enrichment from low-input samples. (**A**) Quantification of localized phosphosites (class 1) across a serial dilution of peptide input; phosphopeptide enrichment purity is shown in purple (top dots; legend on right axis). Each sample was enriched individually. (**B**) Number of quantified phosphopeptides of a serial dilution of naïve CD4 T cells prior to sample processing. (**C**) Coefficient of variation of phosphopeptide quantification across replicates from diluted naïve CD4 T cells. (**D**) Pearson correlation of phosphopeptide intensities across experimental replicates. (**E**) Analysis of linearity between cell input quantity and the intensity of quantified phosphosites. Inset: linear behavior of a representative phosphosite as a function of cell input amount. Outset: histogram of all linearity measurements (Pearson). (**F**) Heatmap depicting kinase activity inference and corresponding substrate quantification across the titration of cell inputs. Individual kinases are represented by individual rows. KS: the absolute number of kinase substrates (KS) for each kinase inferred from the dataset. For each cell input quantity, kinase substrate number is depicted as a color gradient, normalized to the highest input condition. Relative kinase substrate numbers for selected kinases are highlighted in accompanying radar plots; the pie size reflects relative substrate number. n = 4-5 for Jurkat lysate experiment (A), n = 6 for naïve CD4 T cells (B-F).

To benchmark performance in low-input primary T cell samples, we also applied *simplePhos* to a dilution series of primary human naïve CD4 T cells in 96-well format. From as few as 50,000 cells per well, we consistently quantified >5,000 localized phosphosites, and up to ∼15,000 phosphosites with higher cell input (Figure 2B–D). Phosphopeptide enrichment remained >90% across inputs, with strong reproducibility and low variability. Importantly, phosphosite quantification scaled linearly for phospho-serine, threonine, and tyrosine residues (Figure 2E). To assess the biological fidelity of low-input data, we performed kinase–substrate enrichment analysis. Although lower inputs reduced the number of substrates detected, coverage across all major kinase families was preserved, supporting the retention of key signaling information even when performing experiments with limited sample input (Figure 2F). While the subsequent phosphoproteomics analysis in this study did not entail sample input below 30 μg, the ability to perform phosphoproteomics on lower input using the *simplePhos* protocol has proved extremely useful for other experimental studies, particularly for analyzing precious primary samples^40^.

In summary, *simplePhos* enables rapid, scalable phosphoproteomic profiling from low-input samples with high accuracy and reproducibility. This protocol enables analysis across the numerous treatments, timepoints, genetic perturbations, and replicates required for systematic signaling pathway mapping. *SimplePhos* is fully compatible with multichannel pipetting and standard lab equipment, requiring less than one hour from peptide resuspension to MS loading, even when processing full 96-well plates.

### *SimplePhos* enables longitudinal phosphoproteomics of cytokine signaling dynamics

Signal transduction converts extracellular cues into coordinated intracellular responses, with proteins serving as central mediators frequently regulated by post-translational modifications. Phosphorylation, in particular, provides a rapid, reversible, and site-specific mechanism for dynamic control of signaling and plays a central role in immune regulation.

To assess the sensitivity and depth of our *simplePhos* workflow in capturing phosphorylation dynamics, we performed time-resolved phosphoproteomic profiling of primary human CD4 T cells following IFN-β stimulation (Figure 3A). Across all time points, we consistently quantified >13,500 localized phosphosites per sample with >96% enrichment purity ( Supplementary Figure 3A-B). Early phosphorylation patterns were similar between 100 U/mL and 1000 U/mL IFN-β, with both doses eliciting robust and reproducible changes, including significant induction of well-characterized tyrosine phosphorylation sites across the STAT protein family 5 minutes post stimulus (Figure 3B)^41, 42, 43^.

**Figure 3:**
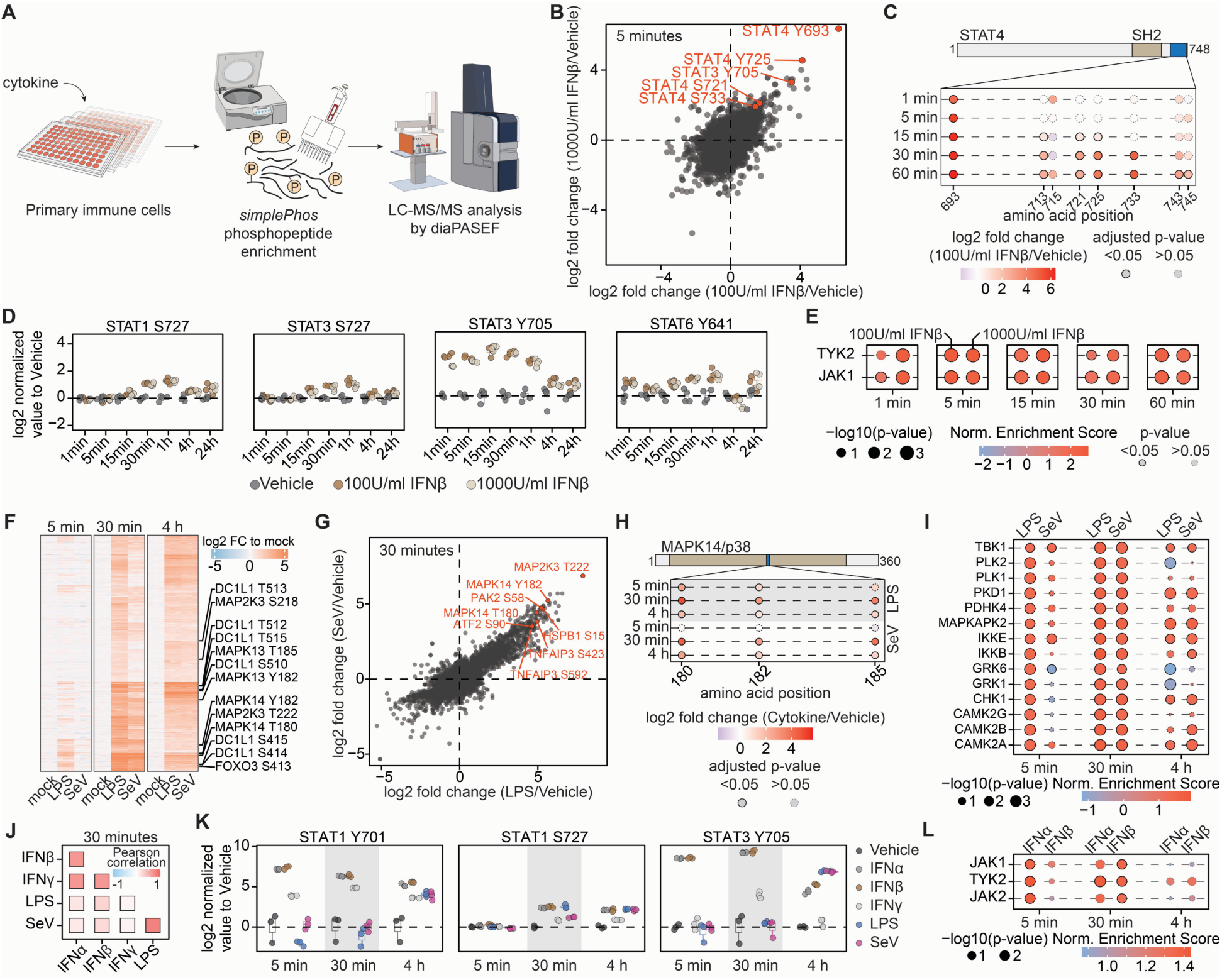
*simplePhos* enables high-resolution temporal phosphoproteomic profiling of cytokine signaling in primary immune cells. (**A**) Experimental workflow used to investigate phosphorylation dynamics in primary immune cells upon cytokine stimulation, employing the *simplePhos* protocol. (**B**) Scatter plot illustrating the regulation of phosphorylation sites in CD4 T cells treated with 100 or 1000 U/ml interferon-β (IFN-β). (**C**) Domain structure of STAT4 with mapped phosphorylation sites identified in samples treated with 100 U/ml IFN-β. (**D**) Time-course boxplots showing phosphorylation kinetics of selected canonical interferon signaling proteins. Each point represents an individual proteomics replicate across the treatment duration. (**E**) Time-resolved kinase activity inference, depicting dynamic changes in activity of key kinases following IFN-β stimulation at two concentrations. (**F**) Heatmap of differentially regulated phosphorylation sites in MDMs following LPS or SeV stimulation. Key signaling proteins and their corresponding phosphorylation sites are highlighted. (**G**) Scatter plot comparing phosphosite abundance in MDMs treated with LPS or SeV for 30 minutes. (**H**) Schematic representation of MAPK14/p38, highlighting quantified phosphorylation sites upon treatment with different cytokines in MDMs. (**I**) Time-resolved kinase activity inference, depicting dynamic changes in activity of key kinases following LPS and SeV treatment in MDMs. (**J**) Assessment of phosphosite similarity between different treatments 30 minutes after stimulation. (**K**) Boxplots showing differential regulation of STAT1 and STAT3 phosphorylation sites following cytokine treatment in MDMs. (**L**) Time-resolved inference of kinase activity in MDMs following IFN-α and IFN-β treatment. For entire figure: n = 4 from two donors for T cell data, n = 3 from one donor for MDM data.

Temporal resolution enabled monitoring of site-specific phosphorylation dynamics. For instance, STAT4 Y693 was rapidly induced within 1 minute post-stimulation, consistent with previous reports studying STAT4 phosphorylation timing following IL-12 stimulation (Figure 3C)^44^. STAT4 Y693 phosphorylation has an essential role in promoting STAT4 dimerization, nuclear translocation, and DNA binding to initiate transcription, and mutation of Y693F abolishes its transcriptional activity^45^. In contrast to the rapid phosphorylation of Y693, STAT4 S721 phosphorylation occurred later – this site is known to modulate transcriptional output under specific conditions^45^. Notably, STAT4 phosphorylation is dispensable for ISG induction; however, activation of STAT4 by IFN-α, IFN-β, or interleukin-12 is critical for IFN-γ production during viral infections^46^.

We also see divergent kinetics of STAT3 phosphorylation, with Y705 phosphorylated more quickly than S727 accompanying IFN-β stimulus (Figure 3D). This is consistent with previous data, as STAT3 Y705 phosphorylation drives dimerization and nuclear translocation – STAT3^Y705F/+^ mice exhibit a profoundly diminished antiviral response - whereas STAT3 S727 phosphorylation contributes to maximal transcriptional output^41, 47^. Other STAT family members also displayed dynamic phospho-regulation (Figure 3D, Supplementary Figure 3C). While STAT1 p-Y701 was not detected in this dataset, delayed induction of p-S727, required for full transcriptional activity, corresponds with ISGF3 complex formation^48^. We also observed temporal dynamics of STAT6 Y641 phosphorylation comparable to that of STAT1 and STAT3 S727. While STAT6 Y641 phosphorylation mediates nuclear translocation and thereby transcriptional activity, its functional relevance is most pronounced in IL-4– and IL-13–mediated signaling^49, 50^. Overall, kinetics of STAT phosphorylation demonstrate the capacity of *simplePhos* to resolve temporally distinct phosphorylation events.

To identify upstream drivers of these phosphorylation dynamics, we applied kinase– substrate enrichment analysis (KSEA) to our time-resolved dataset^51, 52^. We detected rapid activation of JAK1 and TYK2, canonical mediators of IFN-I signaling, within one minute of stimulation (Figure 3E, Supplementary Figure 3D), consistent with their recruitment by the IFN receptor complex. These early events mark the onset of JAK-STAT signaling and precede downstream transcriptional responses. Notably, we also inferred increased activity of DDR1, a receptor tyrosine kinase whose predicted substrate profiles overlap with JAK1/TYK2. While this complicates kinase attribution, it may also reflect functional convergence or crosstalk. Notably, DDR1 has been shown to phosphorylate STAT3 Y705 in inflammatory contexts, including fibrosis, raising the possibility of context-specific modulation of inflammatory pathways^53^. DDR1 expression is largely absent in T cells, underscoring that KSEA results should be interpreted in a cell–type–specific context^54^.

### Dynamic phosphorylation signaling in monocyte-derived macrophages upon innate immune stimuli

To further benchmark *simplePhos*, we next studied phosphorylation changes of monocyte-derived macrophages (MDMs) in response to innate immune stimuli (Figure 3F-3L, Supplementary Figure 3E-F). As key sentinels of innate immunity, MDMs produce and respond to interferons and pro-inflammatory cytokines^55^. Here, we expanded the analysis to include type I interferons (IFN-α and IFN-β), type II interferon (IFN-γ), and the prototypic bacterial and viral innate immune stimuli lipopolysaccharide (LPS) and Sendai virus (SeV), respectively. This design enabled a broader benchmarking of the *simplePhos* pipeline, providing a comprehensive view of stimulus-specific and shared signaling dynamics in primary human innate immune cells.

We first assessed phosphorylation changes after LPS and SeV stimulation (Figure 3F), revealing a pronounced increase in phosphorylation of proteins associated with key signaling pathways activated by LPS as early as 5 minutes after exposure, including Toll-like receptor (TLR) signaling, MAPK cascades, and EGF/EGFR pathways, highlighting the coordinated engagement of multiple signaling modules in the early response to LPS^56, 57^. SeV treatment induced only a limited number of differentially regulated phosphosites at 5 minutes post-stimulation, likely due to delayed kinetics required for SeV infection. By 30 minutes, the SeV phosphosignaling response largely mirrored that of LPS. Both LPS and SeV elicited significant induction of well-characterized phosphorylation sites on proteins such as MAP2K3, PAK2, ATF2, HSPB1, TNFAIP3 (A20), and MAPK14^58^ (Figure 3G). These findings indicate that, although extracellular LPS mainly engages TLR4 and SeV activates RIG-I, their downstream cascades converge on a largely overlapping signaling network. For example, MAPK14 represents a node within this convergence, as it is a well-established component of the canonical pro-inflammatory signaling cascade triggered by LPS, and RIG-I also induces the MAPK14 pathway ^58, 59^. We observed a clear time-dependent increase in phosphorylation at MAPK14 T180 and Y182, its two critical activating residues, as early as 5 minutes following LPS exposure, with maximal phosphorylation reached at 30 minutes – these sites are also phosphorylated upon SeV infection (Figure 3H).

We also performed kinase substrate enrichment analysis to infer upstream kinase activity. In MDMs stimulated with LPS or SeV, this analysis revealed activity of multiple serine/threonine kinases, notably the IKK members TBK1 and IKKε (Figure 3I)^56, 60^. As noted above, LPS appears to drive more rapid signaling, however even at the 5-minute timepoint, we see SeV-mediated activation of IKKε and TBK1, which are downstream of RIG-I and MAVS^61^. These findings indicate that although both LPS and SeV converge on overlapping MAPK-dependent pathways, they do so with distinct temporal dynamics.

We also studied signaling dynamics induced by type-I (IFN-α and IFN-β) and II (IFN-γ) interferons in MDMs. Correlation analysis across different time points (Figure 3J, Supplementary Figure 3G) revealed a strong similarity in phosphorylation signatures across type I and type II interferons, distinct from the correlating signatures of LPS and SeV-induced signaling, highlighting the distinct signaling programs driven by these inflammatory stimuli. Indeed, analyzing JAK–STAT signaling in MDMs, we observe phosphorylation of canonical STAT residues upon treatment with type I and type II interferons. Phosphorylation of STAT1 at Y701, a prototypical activation site of interferon signaling, was rapidly induced by all interferons, with robust increases detectable within 5 minutes and sustained for at least 4 hours^62^. In contrast, LPS and SeV stimulation led to detectable STAT1 Y701 phosphorylation only at 4 hours, likely reflecting secondary signaling events rather than direct pathway activation (Figure 3K). We also observed a delayed increase in STAT1 S727 phosphorylation, consistent with patterns previously noted in CD4 T cells (see Figure 3D). Further, STAT3 Y705 phosphorylation displayed a response broadly comparable to that of STAT1 Y701; however, at early time points, STAT3 activation was markedly more selective for type I interferons, indicating differential sensitivity of STAT family members to distinct cytokine stimuli. We next examined the effects of type I and type II interferon treatment on MDMs. Kinase enrichment analysis of IFN-α– and IFN-β–treated samples revealed a pronounced increase in JAK1 and TYK2 activity, consistent with results previously observed in CD4 T cells (Figure 3E, L). In contrast, kinase activity could not be reliably quantified in IFN-γ–treated samples, likely due to the modest phosphorylation changes observed, suggesting limited pathway activation in this stimulus. While we noted a slightly higher inferred kinase activity in response to IFN-α compared to IFN-β at earlier time points, this difference may reflect a dose-dependent effect.

In summary, our optimized *simplePhos* workflow provides a robust, scalable platform for high-throughput phosphoproteomic profiling in primary human immune cells, even from limited input material. The streamlined sample preparation protocol enables consistent quantification of thousands of high-confidence phosphorylation events per sample. Notably, *simplePhos* effectively resolves dynamic and stimulus-specific phosphorylation patterns of diverse canonical signaling pathways, underscoring its utility for systematic mapping of cell signaling networks.

### CRISPR knockouts in primary T cells enable the attenuation of ISG pathways

Mapping key regulatory nodes within signaling cascades is critical for decoding immune signaling networks and identifying potential therapeutic targets. When available, small-molecule inhibitors can enable pharmacological modulation of signaling nodes, but often suffer from off-target and dose-dependent effects. In contrast, CRISPR-Cas9–mediated gene knockout (CRISPR-KO) provides a precise, genetic approach to dissect the contribution of individual signaling components.

To systematically interrogate the architecture of interferon signaling in primary CD4 T cells, we performed CRISPR-KO of a curated panel of canonical pathway regulators (Figure 4A). Most targeted genes were successfully disrupted without overt effects on cell viability ( Supplementary Figure 4A-D). However, loss of JAK1 and JAK3 led to a pronounced proliferative defect under CD3/CD28 stimulation, consistent with their established roles in IL-2 signaling^63^. Consequently, JAK1 and JAK3 knockout samples were excluded from further quantitative comparisons.

**Figure 4:**
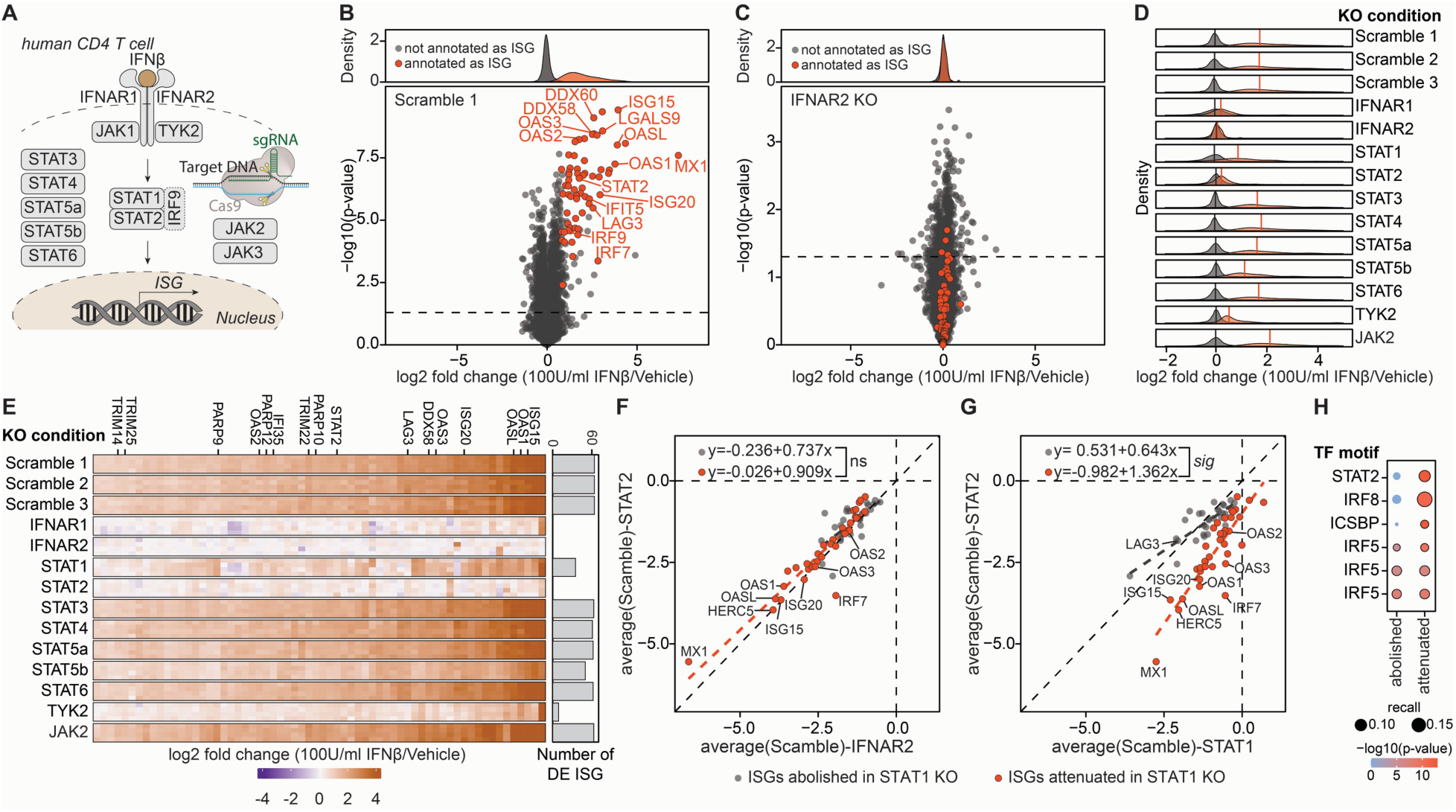
CRISPR knockout in primary CD4 T cells combined with global proteomics identifies key regulators of the interferon response. (**A**) Schematic overview of the type I interferon-β signaling cascade. All CRISPR knockout targets used in this study are indicated. (**B**) Volcano plot showing differential protein abundance in CD4 T cells 24 hours after treatment with 100 U/mL IFN-β compared to vehicle control in scrambled control cells. IFN-stimulated proteins (denoted as ISGs) consistently upregulated across three independent scramble treatments are highlighted in red (adj p-value < 0.05 (Benjamini-Hochberg), log_2_FC > 0.75). (**C**) The same analysis as in (B) was performed in IFNAR2 knockout CD4 T cells, showing reduced ISG response. (**D**) Density distribution highlights ISG-annotated proteins upregulated by IFN-β treatment for all KO conditions. Vertical lines denote mean fold change. (**E**) Quantification of significantly upregulated ISG proteins across different knockout conditions. (**F**) Scatter plot comparing relative protein abundance of ISGs identified by proteomics in IFNAR2 and STAT2 knockout CD4 T cells treated with IFN-β. Linear regression lines are shown, illustrating the degree of similarity in ISG regulation between the two knockout conditions. **(G)** Same as in (F) but comparing STAT1 and STAT2 KOs. ISGs abolished by STAT1 KO highlighted in grey, ISGs still induced / attenuated in STAT1 KO highlighted in red (**H**) Transcription factor motif enrichment analysis for ISGs dependent on STAT1. n = 4, from two donors.

We first assessed the functional impact of ISG expression by knocking out the two subunits of the type I interferon receptor heterodimer: IFNAR1 and IFNAR2. As expected, disruption of either subunit resulted in a substantial reduction in ISG protein expression, confirming both the critical role of the receptor subunits in interferon signaling and the effectiveness of our CRISPR-based genome editing strategy in primary T cells (Figure 4B-C). Knockout of TYK2, a kinase essential for JAK-STAT signaling downstream of type-I interferon, abrogated ISG expression, mirroring the effects of IFNAR1/2 knockout and underscoring its essential role in this pathway (Figure 4D-E).

Encouraged by this, we next evaluated whether knockout of downstream STAT proteins could yield similar suppressive effects on the interferon response. To systematically identify key regulators, we defined a global proteomic ISG signature based on proteins consistently upregulated across three independent scrambled control samples treated with IFN-β (Figure 4D-E, Supplementary Figure 4E). This empirically derived protein set provides a robust and unbiased signature to identify knockouts that modulate interferon-related signaling, enabling a refined exploration of pathway dependencies beyond canonical annotations (Figure 4F). Canonical ISGF3 signaling is mediated by STAT1, STAT2, and IRF9; however, alternative homo- and heterodimeric STAT complexes are also known to regulate ISG expression^64^. To systematically dissect the mechanisms underlying IFN-β-induced ISG expression, we leveraged our knockout dataset to determine whether it can recapitulate non-canonical STAT-driven regulatory circuits, thereby providing an unbiased framework to map the diversity of ISG-inducing complexes. Indeed, we observed a pronounced difference in ISG expression between STAT1 and STAT2 knockout conditions, despite both being components of the ISGF3 complex, alongside IRF9, that mediates ISG expression (Figure 4E). Our data reveals a substantially greater attenuation of ISG expression in STAT2-deficient cells, and this effect was not attributable to differences in gene editing efficiency, as both knockouts demonstrated comparable editing rates ( Supplementary Figure 4A). To further explore this divergence, we compared ISG induction in STAT1, STAT2 or IFNAR2 knockout cells (Figure 4F-G). This revealed a bimodal distribution: while some ISGs (e.g., TRIM22, EPSTI1) were reduced in both STAT1 and STAT2 knockouts, others (e.g., RIG-I, OAS2, IRF7) remained largely unchanged in the STAT1 knockout but were strongly downregulated in STAT2-deficient cells. Indeed, previous studies have shown that STAT1 knockout only partially impairs the antiviral effects of IFN-α, whereas STAT2 knockout leads to a more pronounced loss of interferon-mediated protection against viral infection due to its crucial interaction with IRF9^65^, providing a mechanistic basis for the partial induction of ISGs in STAT1-deficient cells. These results underscore the complexity of signaling networks, whereby canonical components are not always entirely essential for functionality. Indeed, transcription factor motif enrichment analysis of ISGs that were abolished or attenuated in STAT1-KO cells revealed an enrichment for STAT2-binding motifs. Combined with our STAT2-KO phenotype, this indicates that in the absence of STAT1, STAT2 likely mediates a subset of ISG expression (Figure 4H).

Integrating saturating genome editing of key signaling components with global proteomics offers a powerful strategy for systematically dissecting immune signaling pathways. Disrupting core elements of the interferon cascade results in a pronounced loss of ISG expression, highlighting essential components for signal transduction. Notably, we found that knockout of STAT1 versus STAT2 leads to distinct effects on ISG expression, despite their known function as a heterodimer, highlighting asymmetry in their contributions to ISG induction by IFN-β. While global proteomic profiling captures the broad impact of these perturbations on protein abundance, coupling this with phosphoproteomic analysis provides an additional mechanistic dimension.

### CRISPR-KO in CD4 T cells combined with phosphoproteomics enables phospho-pathway mapping

Saturating CRISPR knockout coupled with global proteomics enables systematic mapping of causal links between signaling nodes and ISG expression. However, this approach primarily captures endpoint effects and lacks temporal resolution of signaling dynamics. To address this, we applied our streamlined phosphoproteomics workflow, *simplePhos*, to profile phosphorylation events following short-term IFN-β stimulation (5 and 30 minutes), enabling high-resolution mapping of signaling events ( Supplementary Figure 5A-B).

We first profiled canonical phosphorylation events on STAT family members (Figure 5A–C, Supplementary Figure 5C). STAT1 Y701 phosphorylation was rapidly induced within 5 minutes of stimulation, consistent with its established role in initiating early interferon signal transduction^62^. By contrast, phosphorylation of STAT1 at S727 was delayed, becoming prominent only after 30 min, consistent with previous models showing S727 phosphorylation following chromatin association^48^. In IFNAR1, IFNAR2, and TYK2 knockout cells, STAT1 phosphorylation was completely abolished, underscoring the essential role of the IFNAR1/2 receptor and TYK2 receptor-associated kinase in pathway initiation. Phosphorylation of STAT3 Y705 and STAT4 Y693 was also absent in IFNAR1/2 and TYK2 knockouts.

**Figure 5:**
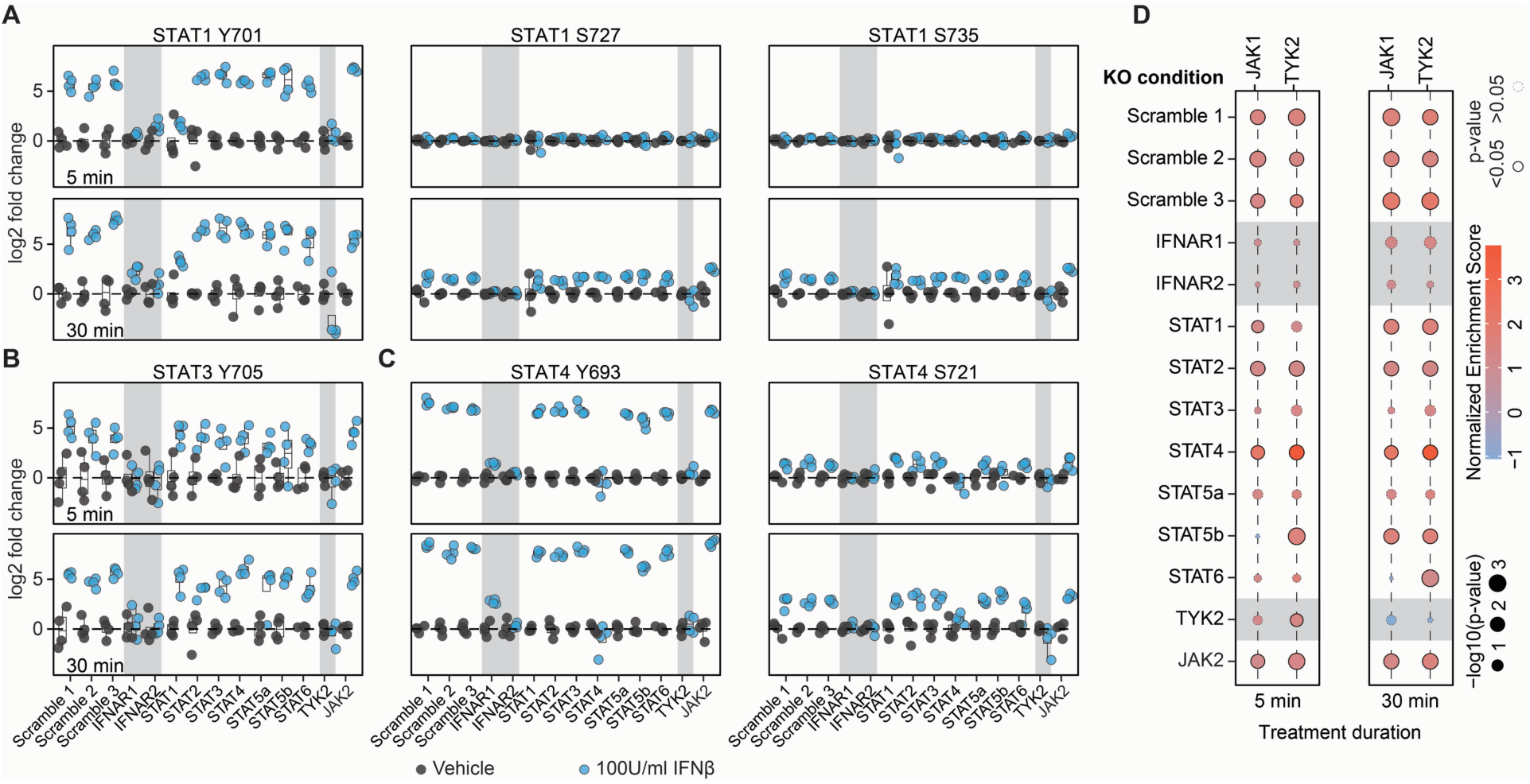
simplePhos enables phosphoproteomic profiling in CRISPR knockout primary CD4 T cells. (**A**–**C**) Quantitative analysis of canonical phosphorylation sites on STAT1 Y701, S727, S735, STAT3 Y705, and STAT4 Y693 and S721 in CRISPR knockout CD4 T cells following interferon-β stimulation for 5 and 30 minutes. Phosphorylation levels are shown relative to vehicle-treated controls with matching genetic backgrounds. Knockouts causing significantly altered downstream phosphorylation relative to scrambled controls are indicated with a grey background. (**D**) Kinase-substrate enrichment analysis of phosphosites significantly regulated by 100 U/ml IFN-β at 5 and 30 minutes. n = 4, from two donors.

To complement these phosphorylation profiles, we conducted kinase substrate enrichment analysis to infer upstream signaling activity across the various knockout conditions (Figure 5D). As anticipated, loss of IFNAR1, IFNAR2, and TYK2 abolished predicted JAK1 and TYK2 activity, consistent with their role as receptor-proximal kinases that initiate type I interferon signaling. In contrast, STAT1 and STAT2 knockouts had minimal effect on JAK1 or TYK2 activity, in line with their position downstream of these kinases in the signaling cascade^62^. Minor changes in JAK1 and TYK2 activity were detected in other STAT knockout conditions, but in-depth analysis of kinase-substrate recognition sites indicates that these effects are likely due to substrate loss from those STAT knockouts rather than alterations in general IFN signaling. Overall, these results not only validate our analytical approach but also reinforce the hierarchical architecture of the IFNAR–JAK–STAT axis.

Together, this integrated approach, combining CRISPR-based perturbations with time-resolved phosphoproteomics, offers a powerful framework for dissecting cytokine signaling architecture in primary immune cells. It enables precise mapping of phosphorylation-dependent events, inference of kinase activity states, and systematic identification of signaling disruptions upon genetic perturbation.

### Base editing enables precise interrogation of phosphorylation sites in primary CD4 T cells

To complement gene knockout approaches and dissect the functional relevance of individual phosphosites, we employed adenine base editing (ABE) to introduce targeted single-nucleotide substitutions at key tyrosine residues within the interferon signaling cascade (Figure 6A)^9^. Focusing on critical tyrosine residues in the canonical interferon signaling cascade: STAT1 Y701, STAT2 Y690, and STAT3 Y705, we optimized an electroporation protocol in activated primary CD4 T cells to co-transfect ABE8e mRNA and sgRNA targeting these tyrosine sites, achieving editing efficiencies up to 85% with minimal off-target effects for STAT1 Y701, and up to 97% editing efficiency targeting STAT3 Y705 ( Supplementary Figure 6A). As is frequently observed with base editing, efficiency varied in a locus-dependent manner^66^. In particular, suboptimal modification at the targeted STAT2 Y690 precluded robust functional assessment of this phosphosite ( Supplementary Figure 6A).

**Figure 6:**
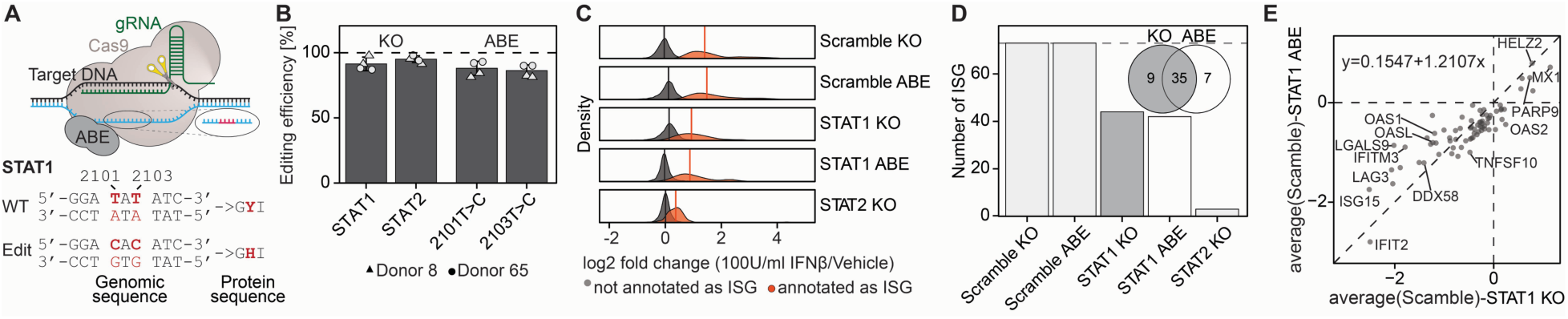
Base editing in primary CD4 T cells enables precise perturbation of phosphorylation sites. (**A**) Base editing strategy introducing the STAT1 Y701H mutation. (**B**) Editing efficiency of STAT1 and STAT2 knockout samples, as well as STAT1 Y701H base-edited samples, across two independent donors. (**C**) Density distribution of ISG-annotated versus non-ISG proteins across all conditions. Vertical lines represent mean fold changes. ISGs were empirically defined based on consistent induction in scrambled control samples (adj p-value < 0.05, log_2_FC > 0.75). (**D**) Number of significantly upregulated ISG proteins identified per genetic perturbation. Venn diagram showing the overlap of ISGs that are abolished upon STAT1 knockout or STAT1 Y701H base editing. (**E**) Scatter plot comparing relative protein expression of ISGs between STAT1 knockout and STAT1 Y701H base editing samples after IFN-β treatment. A linear regression line is shown to assess correlation and the similarity in transcriptional control by full STAT1 loss versus loss of Y701 phosphorylation. n = 6, from two donors.

Next, following IFN-β stimulation for 24 hours, we performed global proteomic profiling of base-edited cells alongside corresponding CRISPR-Cas9 STAT knockouts and scrambled controls (Figure 6B-E, Supplementary Figure 6B-E). While CRISPR-mediated knockout of STAT1 led to near-complete loss of the encoded protein, base editing of STAT1 Y701 preserved STAT1 protein levels, indicating that phosphosite editing perturbs signaling without inducing protein degradation (Figure 6B, Supplementary Figure 6B-C). Quantitative analysis of ISG products revealed that STAT1 Y701 editing markedly suppressed ISG induction, closely mirroring the ISG global proteomic profiles of STAT1 knockout cells (Figure 6D-E, Supplementary Figure 6E). These findings demonstrate that disrupting the single phosphorylation site Y701 on STAT1 is sufficient to impair interferon signaling, underscoring the mechanistic precision and utility of base editing for probing phosphorylation-dependent pathways in primary immune cells. Notably, STAT1 knockout and STAT1 Y701H-edited cells showed comparable residual ISG expression profiles with substantial overlap (Figure 6C, D).

Collectively, these findings establish base editing as a precise and scalable approach for probing the signaling consequences of specific post-translational modifications in primary human immune cells. Disruption of STAT1 Y701 alone was sufficient to phenocopy the effects of full STAT1 knockout, underscoring the critical role of this phosphorylation event in mediating interferon signaling. This strategy enables high-resolution mechanistic dissection of signaling outcomes in their native cellular context without altering overall gene and protein integrity.

## Discussion

Deciphering signal transduction in primary immune cells is essential for understanding the molecular networks mediating inflammation, host defense, and immune dysregulation^67^. Fine-grained, time-resolved mapping of these pathways reveals how immune cells interpret extracellular cues, such as cytokines or pathogen-associated molecular patterns, and identifies actionable nodes for therapeutic intervention. However, capturing these dynamic and transient signaling events remains technically challenging due to limited quantity of primary samples and the rapid kinetics of post-translational modifications. This study presents a blueprint for systematically dissecting these pathways by integrating saturating genetic perturbations with mass spectrometry–based analyses. This combined strategy enables both causal inference, through precise gene perturbation, and broad discovery, through unbiased proteome-wide profiling, thereby providing a powerful framework to unravel signaling architecture.

To enable this study, we developed *simplePhos*, a scalable and sensitive phosphoproteomic workflow, and validated it by interrogating diverse signaling pathways in primary immune cells. *SimplePhos* enables robust and scalable phosphosite enrichment from minimal input material without the need for specialized preparative instrumentation, making it broadly accessible and well-suited for analysis of precious primary samples. Applying this method to study innate immune signaling in stimulated CD4 T cells and monocyte-derived macrophages, we identified canonical phosphorylation events, including STAT1 Y701, STAT3 Y705, and STAT4 Y693 for interferon and MAPK14 T180 and Y182 for LPS and SeV, among others, each exhibiting distinct temporal kinetics. Overall, these findings demonstrate the power of *simplePhos* to resolve dynamic signaling events and uncover their temporal regulation. To move beyond descriptive observations toward mechanistic understanding, we integrated protein abundance and phosphoproteomic profiling with targeted genetic perturbations.

Disruption of core JAK/STAT pathway components such as IFNAR1, IFNAR2, and TYK2 led to attenuation of ISG induction and IFN-induced phosphorylation, thereby revealing functional dependencies of interferon signaling. Furthermore, precise base editing of a single phosphorylation-modified amino acid, STAT1 Y701, fully recapitulated the effects of STAT1 gene knockout. This result confirms this site’s essential role in interferon signaling in primary human T cells, and underscores the precision of base editing as a tool to dissect site-specific phosphorylation functions without the pleiotropic consequences of complete protein loss. Our pathway analysis included 11 gene knockouts and 3 non-targeting controls, each run in quadruplicate and treated with IFN-β across two time points with matched vehicle controls. In total this yielded 224 phosphoproteomic samples, and this scale of global phosphoanalysis was enabled by the *SimplePhos* protocol.

While crucial for immune function and regulation, phosphorylation is only one of many signaling modalities, and focusing on a single PTM at a time inevitably represents a simplification of the underlying biological complexity^68, 69, 70, 71^. Indeed, future work will include orthogonal PTM enrichment strategies to capture a broader range of regulatory modifications beyond phosphorylation in the same sample^72, 73, 74^. With continued development of top-down proteomics, eventually one could envision analyzing PTMs at the intact proteoform level with high throughput^75, 76^. Additionally, continued development of cross-linking mass spectrometry workflows may eventually enable interrogation of protein-protein interaction dynamics. These emerging approaches may one day provide unprecedented systematic insights of biochemical dynamics, enabling a more comprehensive understanding of signaling and regulatory mechanisms.

While loss- or gain-of-function animal models exist for many immune-relevant genes, their generation and maintenance are often labor-intensive, time-consuming, and expensive^77^. Such models are often generated in murine systems, which only partially recapitulate human immune biology. In contrast, saturating CRISPR/Cas9 knockout or base editing offers an attractive alternative, enabling the efficient generation of cells with precise genetic perturbations directly in human immune cells, and across diverse genetic backgrounds. This approach accelerates functional studies and enhances the translational relevance of the findings. That said, there are abundant opportunities for improvement of genetic engineering approaches. This is particularly relevant for base editing, where efficiency is highly locus-specific^78^. Consistent with this, we observed residual ISG induction in incompletely edited samples, underscoring the need for careful genotypic validation of editing outcomes. Ongoing advances in gene editing are poised to further boost efficiency and precision, opening exciting new opportunities for pathway mapping in primary cells ^79^

Cellular signaling is highly complex and context-dependent. For example, although phosphorylation of STAT3, 4 and 6 are induced by interferon-β, knockout of these proteins did not attenuate ISG expression. This observation highlights the inherent complexity of signaling networks, where phosphorylation on specific transcription factors may occur without directly contributing to canonical transcriptional outputs. Instead, such modifications may reflect cross-talk with parallel cytokine pathways, context-specific regulation, and serve as priming events for alternative cellular responses, underscoring the layered, interconnected architecture of cytokine signaling. It is for this reason why observational signaling studies are insufficient to understand the mechanistic underpinnings of signaling outcomes. Integration of saturating genetic perturbations with deep proteomic profiling thus provides a biochemical perspective of signaling causality that was not previously possible, and offers the ability to refine pathway models across diverse cellular contexts.

In summary, our integrative platform combining functional genetics with systematic global and phosphoproteomics demonstrates a powerful framework to dissect dynamic immune signaling with exceptional depth and precision. Beyond advancing fundamental insights into immune regulation, this strategy is broadly applicable for mapping diverse signaling events across immune and non-immune cell types. Critically, introducing targeted genetic interventions enables mechanistic analyses of signaling dynamics and proteomic outcomes directly in human samples, thereby establishing a versatile blueprint for decoding complex signaling circuits. Looking forward, this framework has the potential to refine further our understanding of how cells respond to external stimuli in health and disease.

## Methods

### Primary T cell enrichment and culture

Trima LRS cones from healthy anonymous donors were purchased from LifeSouth. Leuko-enriched blood was mixed 1:1 with PBS (Corning) containing 2 mM EDTA (Invitrogen). Dilute blood was layered on top of Ficoll (Cytiva Life Sciences) and centrifuged for 10 minutes at 1200xg in a SepMate tube (StemCell), buffy coat was washed three times in PBS-EDTA, and a sample of PBMCs was reserved for flow cytometry analysis. CD4 T cells were enriched by negative selection using a CD4 T cell isolation kit (StemCell), and a sample of enriched cells was also reserved for flow cytometry analysis. CD4 T cell enrichment was quantified by labeling with FITC-conjugated anti-CD4 (BD, clone OKT4, 1:100 dilution) and APC-conjugated anti-CD25 (BioLegend, clone BC96, 1:100 dilution), and flow cytometry analysis was performed on a FACS Symphony A3 (BD). T cells were cultured in RPMI (Corning) supplemented with 10% fetal calf serum (Cytiva Life Sciences), 2 mM L-glutamine, 110 μg/mL sodium pyruvate, 110 μg/mL penicillin-streptomycin (Avantor Sciences), 25 mM HEPES (Avantor Sciences), and 40-80 U/mL IL-2 IS (Miltenyi).

### Time course of interferon treatment of CD4 T cells

T cells were stimulated using plate-bound anti-CD3 and suspension anti-CD28 (Tonbo) for 2-3 days, and thereafter were expanded and maintained at a confluence of 1-3 million cells/mL. Cells were treated with interferon beta at 100 or 1000 U/mL (Peprotech), and each time point was accompanied by a time-matched negative control. Following IFN treatment, signaling was arrested by treatment with 4x volume of ice-cold PBS and washed 2-3 times in cold PBS prior to pelleting and snap freezing on dry ice and storage at -80 °C.

### MDM differentiation and treatment

CD14+ peripheral blood monocytes were positively isolated from peripheral blood mononuclear cells (PBMCs) isolated from leukopaks (NY Biologics) from anonymous donor by using CD14 MicroBeads (Miltenyi Biotec, 130-050-201), and differentiated into human monocyte-derived macrophages (MDMs) by culturing in RPMI-1640 (Invitrogen) containing 10% heat-inactivated human AB serum (Sigma) and recombinant human M-CSF (20 ng/ml; Peprotech) for 5–6 days as previously described^80^.

### Macrophage stimulation and collection for proteomic/phospho-proteomic analysis

CD14+ monocytes were plated at ∼3×10^6^ cells per well in 6-well plates and differentiated into macrophages (∼2-2.5×10^6^ cells per well after differentiation). One day prior to innate immune stimulation, the media was removed, and 1.5 mL of fresh RPMI media with 10% FBS was added to the cells. For stimulation, a 4X stock of each stimulus was made up in media, and 500 μL was added to cells for a total volume of 2 mL. Cells were stimulated with 100 IU/mL IFN-β, 1 nM IFN-α2a (PBL assay Science; Cat. #11101-2), 1 nM IFN-γ (PBL assay Science; Cat. #11500-1), 100 ng/mL lipopolysaccharide (LPS) (InvivoGen; Cat. #tlrl-eblps), 15 hemagglutinin assay units/mL Sendai Virus strain Cantell (Charles River Laboratories), or mock-treated (RPMI media with 10% FBS). Cells were collected after 5 min, 30 min, 4 hours, 24 hours, or 36 hours after stimulation. After stimulation, the media was removed, ice-cold 1X PBS (Gibco; Cat. # 14190144) was added to each well, and plates were placed on ice. PBS was removed, plates were washed twice with ice-cold PBS, and 250 μL of lysis buffer (1% sodium deoxycholate powder in 100mM Tris, pH 8.0) containing phosphatase and protease inhibitor was added to each well. Cells were then scraped, lysates were collected into pre-chilled 1.5 mL Protein Lobind Eppendorf tubes, and flash frozen prior to storage at -80°C.

### Proteomic sample preparation

For high input samples (>100 μg per sample): Protein pellets were resuspended in a buffer containing 50 mM Tris (pH 8) (Invitrogen), 1% sodium deoxycholate (RPI), 10 mM TCEP (Thermo Scientific), and 40 mM CAA (TCI Chemicals), followed by heating at 95°C for 10 minutes. Samples in Eppendorf Lobind tubes (optimization experiments) were then sonicated using a Bioruptor (Diagenode), or for other samples (interferon titration CD4 T cells, all experiments on MDMs) genomic DNA was digested using Universal Nuclease (Thermo Fisher Scientific) at a concentration of 10U per 200 μL of sample and incubated at 37°C for 30 minutes. Protein concentration was measured using the BCA assay as per the manufacturer’s protocol (Thermo Fisher Scientific). Next, a maximum of 200 μg of protein was digested overnight at 37°C with LysC/Trypsin (Promega) at a protein-to-enzyme ratio of 1:200. The following day, the samples were acidified with TFA and desalted using HLB plates (Waters). After elution, the peptide concentration was assessed using the Quantitative Colorimetric Peptide Assays (Thermo Fisher Scientific). For global proteomics analysis, approximately 100 ng of clean peptides were loaded onto EvoTips following the manufacturer’s guidelines (EvoSep Biosystems). The rest of the samples were dried to almost completeness in a SpeedVac (LabConco) and stored at -80°C for phosphopeptide enrichment by *simplePhos*.

For low input samples (<100 μg per sample; knockout and naïve T cell samples): Low input samples were harvested in 50 mM Tris (pH 8), 0.2% DDM (Thermo Scientific), 10 mM TCEP, and 40 mM CAA. Genomic DNA was then digested using Universal Nuclease (Thermo Fisher Scientific) at a concentration of 10U per 200 μL of sample and incubated at 37°C for 30 minutes. Protein concentration was measured using the BCA assay as per the manufacturer’s protocol (Thermo Fisher Scientific). Next, proteins were digested overnight at 37°C with LysC/Trypsin (Promega) at a protein-to-enzyme ratio of 1:200. For global proteomics analysis, approximately 100 ng of digested peptides were loaded onto EvoTips following the manufacturer’s guidelines (EvoSep Biosystems). The rest of the samples were dried to almost completeness in a SpeedVac and stored at -80°C for phosphopeptide enrichment by *simplePhos*.

### *SimplePhos* Phosphoproteomic sample preparation

Peptides were resuspended for 10 minutes in a loading buffer containing 80% ACN (Avantor Sciences), 5% TFA (Avantor Sciences), 0.1 M glycolic acid (Thermo Scientific), 0.1% DDM, and centrifuged for 5 minutes at 16,000 xg to remove any insoluble peptides. Simultaneously, Zr-IMAC HP beads (MagResyn, Resyn Biosciences; 5 μL of beads per sample) were washed three times with the same loading buffer. The washed beads were then added to each sample (30 μg of peptides if not otherwise specified), allowing phosphopeptides to be enriched for 10 minutes in a Thermo Mixer (Thermo Fisher Scientific) at 1200 rpm 25°C. The beads were subsequently immobilized using a Magnetic Stand-96 (Thermo Fisher Scientific), washed with loading buffer, then washed in 80% ACN with 1% TFA, 0.1% DDM, and finally washed with 10% ACN with 0.1% TFA, 0.1% DDM. All washing steps were performed on a Thermo Mixer. Phosphopeptides were then eluted from the beads using 80 μL fresh 1% ammonia (Millipore Sigma), 0.1% DDM for 10 minutes shaking 1200 rpm 25°C, then immediately acidified with 20 μL 10% formic acid (Thermo Scientific). Samples were then loaded onto pre-conditioned EvoTips (EvoSep Biosystems) according to the manufacturer’s instructions.

### Mass spectrometry

Samples were injected into an Evosep One system (EvoSep Biosystems) coupled to a timsTOF Pro2 mass-spectrometer (Bruker Daltonics). Samples were analyzed using the 40 SPD Whisper Zoom predefined gradient unless otherwise stated, using a commercial analytical column (15 cm 75 μm CSI, Aurora Elite, IonOpticks). The mass-spectrometer operated in positive polarity for data collection using data-independent acquisition (diaPASEF) mode. DIA isolation windows were optimized to the expected precursor ion density for global and phosphoproteomics samples separately^81^. For the synchoPASEF method, the method was designed with pyDIA employing the same windowing range as for diaPASEF^81^.

### Data analysis

The raw files were processed in Spectronaut^82^ v20 in library-free mode (directDIA). For the database search, we used the UniProt Homo sapiens database (UP000005640, downloaded on 19.8.2023, 20,420 entries). We defined Carbamidomethylation (C) as a fixed modification, and Acetyl (Protein N-term), and Oxidation (M) were specified as variable modifications. For phosphopeptide-enriched samples, we further defined phosphorylation (S, T, Y) as a variable modification and enabled the localization workflow. PTM site collapse was performed in Spectronaut with default settings.

All remaining downstream data analysis was performed in R (version 4.4.0 and newer). For global and phosphoproteomics data, we only considered proteins or phosphosites where, for at least one condition, at least 3 out of 4 replicates (or 2 out of 3 for MDM samples), were quantified. Intensities were normalized by median centering and log_2_-scaling. Missing values were imputed by k-NN imputation (global proteomics) or SLSA imputation (phosphoproteomics) and downshift sampling, for proteins missing at random or not at random, respectively ^83^. Samples were normalized by donor (if applicable), and then fold change was determined relative to the time-matched control samples. All statistical comparisons were performed using two-tailed Student’s t-tests (global proteomics) or limma (phosphoproteomics), with Benjamin-Hochberg p-value correction.

### Cas9 knockouts

Guide RNA (sgRNA) was ordered from Synthego and resuspended to a final concentration of 100 μM in 10 mM Tris, pH 7.4 (Lonza) prior to use. For each RNA nucleofection reaction, 2 μL of sgRNA was mixed with 2 μL of 40 μM Cas9-NLS protein to make 4 μL of Cas9-RNP complex and incubated at 25°C for 15 minutes, then placed on ice prior to electroporation. Cas9-RNP complexes were electroporated into 1,000,000 stimulated primary CD4 T cells using the Lonza Primary P3 nucleofection kit and 4D Nucleofector system using program EO-115. After nucleofection, cells were re-stimulated using T cell activation and expansion beads (Miltenyi) and 80 IU/mL IL-2 IS (Miltenyi), and cells were split every 2-3 days. Knockout efficiency was assayed by ICE analysis (Synthego) of genomic DNA amplicons by Sanger sequencing.

### Cloning of pABE8e(V106W)-SpRY

To generate the pABE8e(V106W)-SpRY (D10A) plasmid, cloning was carried out using isothermal assembly, with primers ordered from Integrated DNA Technologies. Briefly, desired vector components were amplified by PCR using Phusion U Green Hot Start II DNA Polymerase (Thermo Fisher Scientific). ABE8e (TadA-8e V106W) (Addgene Plasmid #138495) and pCAG-CBE4max-SpRY-P2A-EGFP (RTW5133) (Addgene Plasmid #139999). The resulting amplicons were purified using the QIAquick PCR Purification Kit (Qiagen). Purified fragments were assembled with NEBuilder HiFi DNA Assembly Master Mix (New England Biolabs) to create plasmid LpAS3902, and assemblies were transformed into One Shot Mach1 cells (Thermo Fisher Scientific). Transformed cells were plated on LB agar containing 50 μg/mL carbenicillin.

### Cas9 base editor mRNA production

SpRY Cas9n ABE8e V106W mRNA was amplified from plasmid LpAS3902 using previously described primers^84^. Following DpnI treatment and column purification (Takara), linear template DNA was quantified, analyzed by agarose electrophoresis, and used for in-vitro RNA transcription (IVT) using Codex T7 RNA polymerase, CleanCap AG, and moUTP. IVT reactions were treated with DNAse, Antarctic phosphatase (New England Biolabs), and purified using a MEGAClear kit (Invitrogen). Final mRNA was quantified by NanoDrop, size was analyzed by agarose gel electrophoresis, and aliquots were snap frozen on dry ice and stored at -80°C.

### Base editing of primary T cells

For each electroporation, 2 μg Cas9 base editor mRNA was mixed with 2 μL sgRNA. 1 million cells were pelleted and resuspended in 20 μL of Lonza P3 buffer, then cells were mixed with Cas9 mRNA + sgRNA, then immediately transferred to a Nucleofector nanocuvette (Lonza). For optimization experiments, the appropriate nucleofection program was applied, whereas program EN-138 was subsequently used for samples intended for proteomics analysis. Following nucleofection, 80 μL of warm complete RPMI was added to each well, and the nucleofector plate was incubated at 37°C for 30 minutes. Complete RPMI with 160 U/mL IL-2 IS and CD2/3/28 stimulation beads (Miltenyi) was created, and 100 μL was aliquoted out into a U-bottom 96-well plate. After the 30-minute incubation, the cells were resuspended and transferred from the nanocuvette into the 96-well U-bottom. These cells were grown for 2 weeks with the media changed and the cells split as needed for optimal growth.

### Genomic sequencing of edited primary T cells

Cells were washed 2-3 times with ice-cold PBS and banked at -20°C until ready to be processed. 100 μL of QuickExtract (LGC Biosearch) was added to each sample and transferred to a 96-well PCR plate. This was incubated at 65°C for 15 minutes, then inactivated at 98°C. PCR reactions were then set with the appropriate primers using SuperFi polymerase according to the manufacturer’s recommended protocol. The PCR reactions were sent for Sanger sequencing with their respective sequencing primers (Azenta). The Cas9 knockout samples were analyzed using ICE analysis (Synthego) to determine their knockout efficiency. Base editing efficiency in primary T-cells was analyzed using BEAT (Base Editing Analysis Tool)^85^.

### Flow cytometry

Approximately 500,000 cells per condition were washed 2–3 times with ice-cold PBS. For viability assessment, samples were stained with Zombie Violet Fixable Viability Dye (BioLegend) in 100 μL of 1:1000 dilution in PBS, then incubated on ice for 20 minutes. Afterwards, samples were mixed with 100 μL of antibody master mix, prepared in FACS buffer (PBS supplemented with 2% FBS and 2 mM EDTA), and incubated on ice for 30 minutes. Following staining, cells were washed three times with FACS buffer and subsequently fixed in 4% paraformaldehyde (PFA) in PBS. Unstained and single stained controls were included for all flow cytometry analyses. For quantification of absolute cell numbers, 50 μL of CountBright Absolute Counting Beads (Thermo Fisher Scientific) were added directly to the fixed samples. Data acquisition was performed on a BD FACSymphony A3 flow cytometer. Flow cytometry data was analyzed using FlowJo, and cell counts derived using Microsoft Excel.

---

**Supplementary Figure 1:**
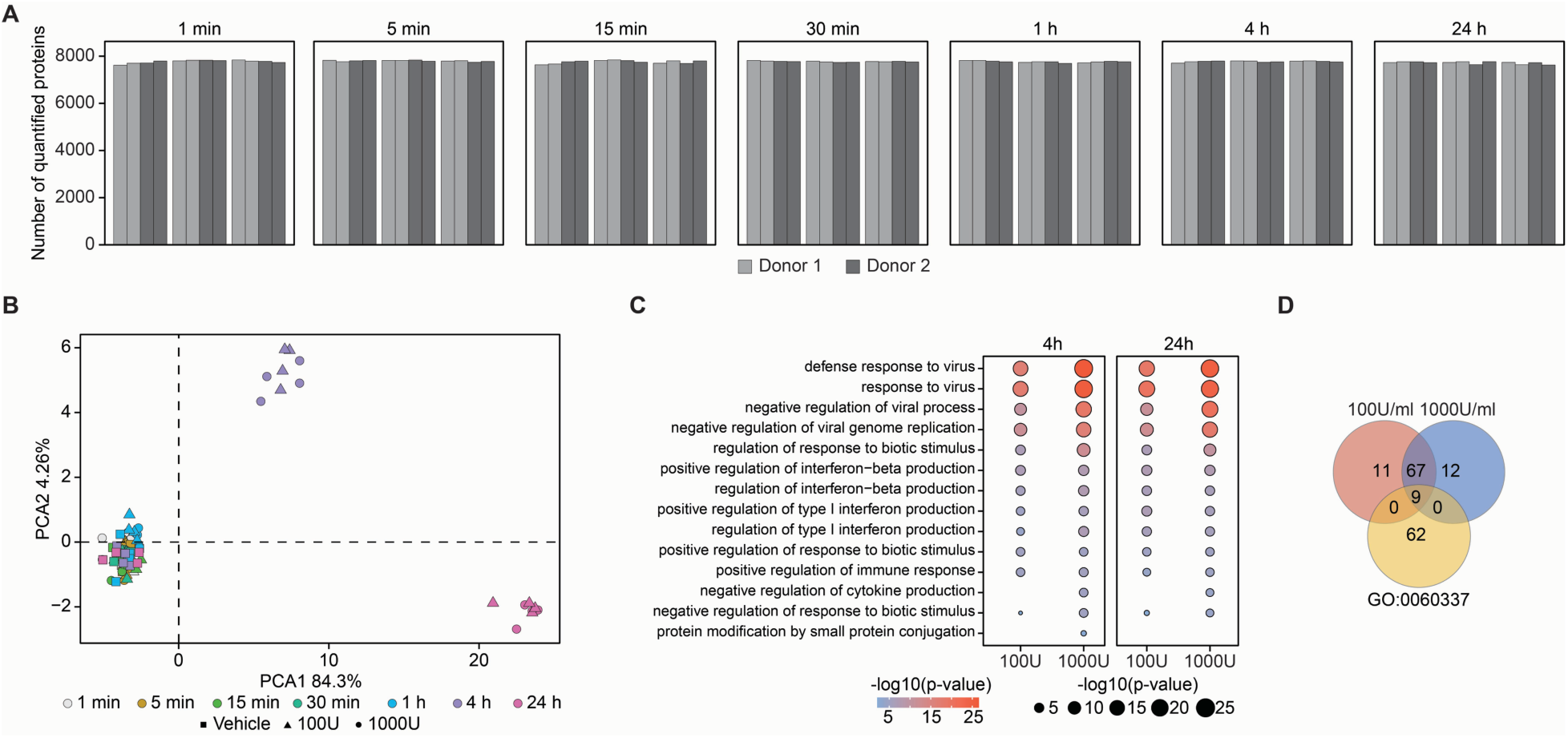
Proteomic Overview of IFN-β–Treated Primary Human CD4 T Cells. (**A**) Number of quantified proteins in each proteomics sample. (**B**) Principal component analysis (PCA) based on significantly regulated proteins following IFN-β stimulation. (**C**) Gene Ontology (GO) enrichment analysis of significantly regulated proteins, highlighting pathways associated with interferon signaling. (**D**) Venn diagram showing shared and unique ISGs across the two treatment conditions, compared with proteins included in the Gene Ontology term GO:0060337.

**Supplementary Figure 2:**
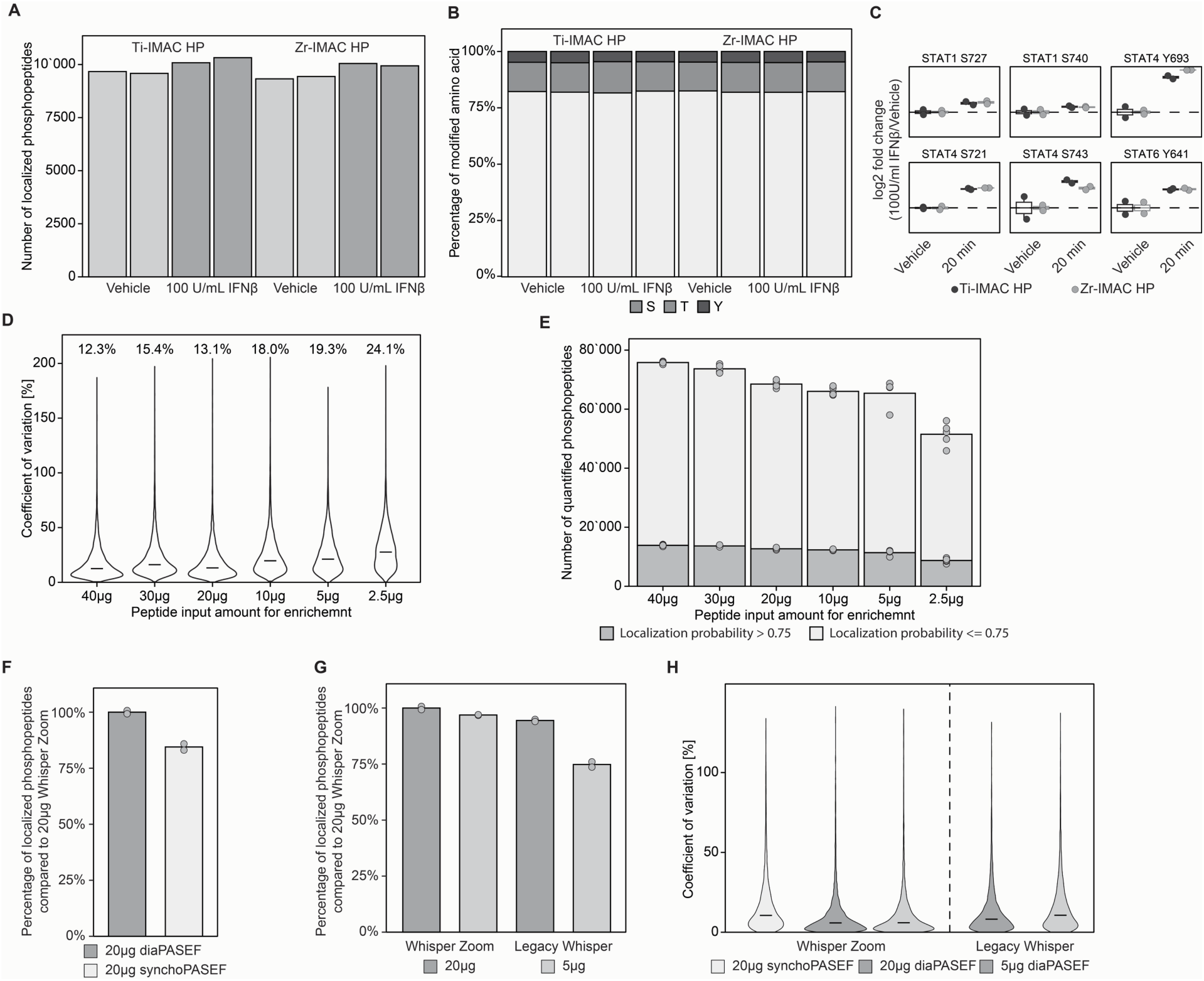
Optimization of the simplePhos workflow for phosphoproteomics in primary immune cells. (**A**) Comparison of the number of class 1 phosphopeptides enriched using Ti-IMAC HP versus Zr-IMAC HP magnetic beads using CD4 T cells. (**B**) Bar plot showing the distribution of class I phosphopeptides by residue type, indicating the percentage of phosphorylation events occurring on serine, threonine, and tyrosine residues. (**C**) Box plots displaying differential phosphorylation of canonical STAT protein sites following interferon-β treatment, using enrichment with Ti-IMAC HP or Zr-IMAC HP beads. (**D**) Coefficient of variation (CV) of phosphopeptide quantification across biological replicates from a peptide input dilution series. (**E**) Quantification of class 1 and non-localized phosphosites across different peptide input levels to evaluate sensitivity and yield of confidently localized sites. (**F**) Number of localized phosphopeptides identified using either diaPASEF or synchro-PASEF acquisition modes on the timsTOF Pro2 platform, using 20 µg peptide input for the simplePhos enrichment, and the EvoSep Whisper Zoom gradient. (**G**) Impact of EvoSep chromatographic separation methods on phosphopeptide quantification using different peptide inputs. **(H)** CV of phosphopeptide quantification across replicates comparing multiple variables: Acquisition strategy (synchro-PASEF vs diaPASEF), EvoSep chromatography separation method (“Whisper Zoom” vs “Legacy Whisper”), and peptide input amounts into the simplePhos enrichment pipeline.

**Supplementary Figure 3:**
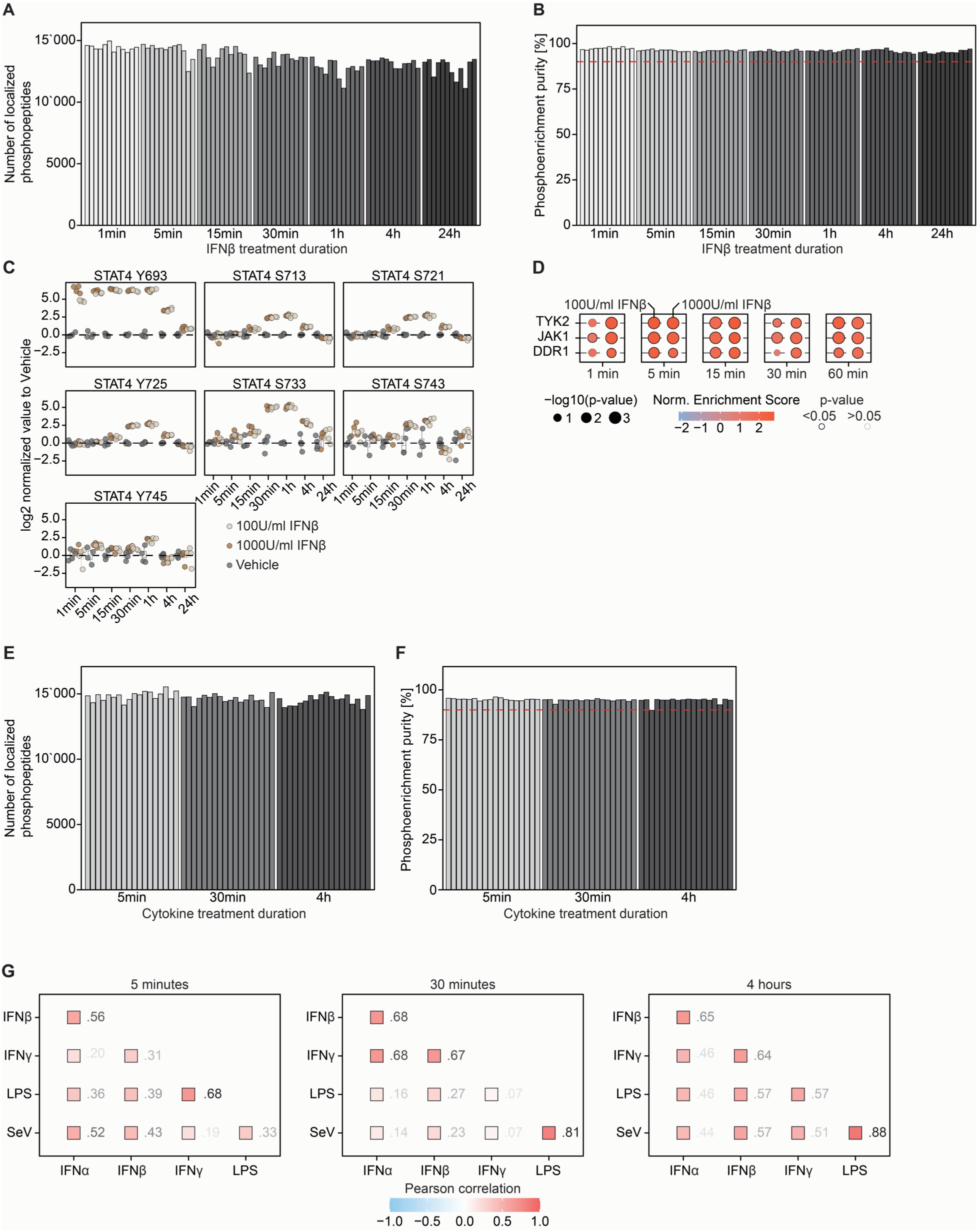
*simplePhos* enables high-resolution temporal phosphoproteomic profiling of cytokine signaling in primary immune cells. (**A**) Number of localized phosphopeptides in each CD4 T cell proteomics sample. (**B**) Purity of phosphopeptides across all samples; 90% threshold of phosphopeptide enrichment purity is shown in red. (**C**) Time-course boxplots showing phosphorylation kinetics of STAT4 phosphorylation sites. (**D**) Time-resolved kinase activity inference, depicting dynamic changes in activity of key kinases following interferon-β stimulation at two concentrations in CD4 T cells. (**E**) Number of localized phosphopeptides in each MDM proteomics sample. (**F**) Purity of phosphopeptides across all MDM samples; 90% threshold of phosphopeptide enrichment purity is shown in red. (**G**) Assessment of signaling similarity by Pearson correlation across phosphosites 5, 30 minutes, and 4 hours after cytokine stimulation of MDM samples.

**Supplementary Figure 4:**
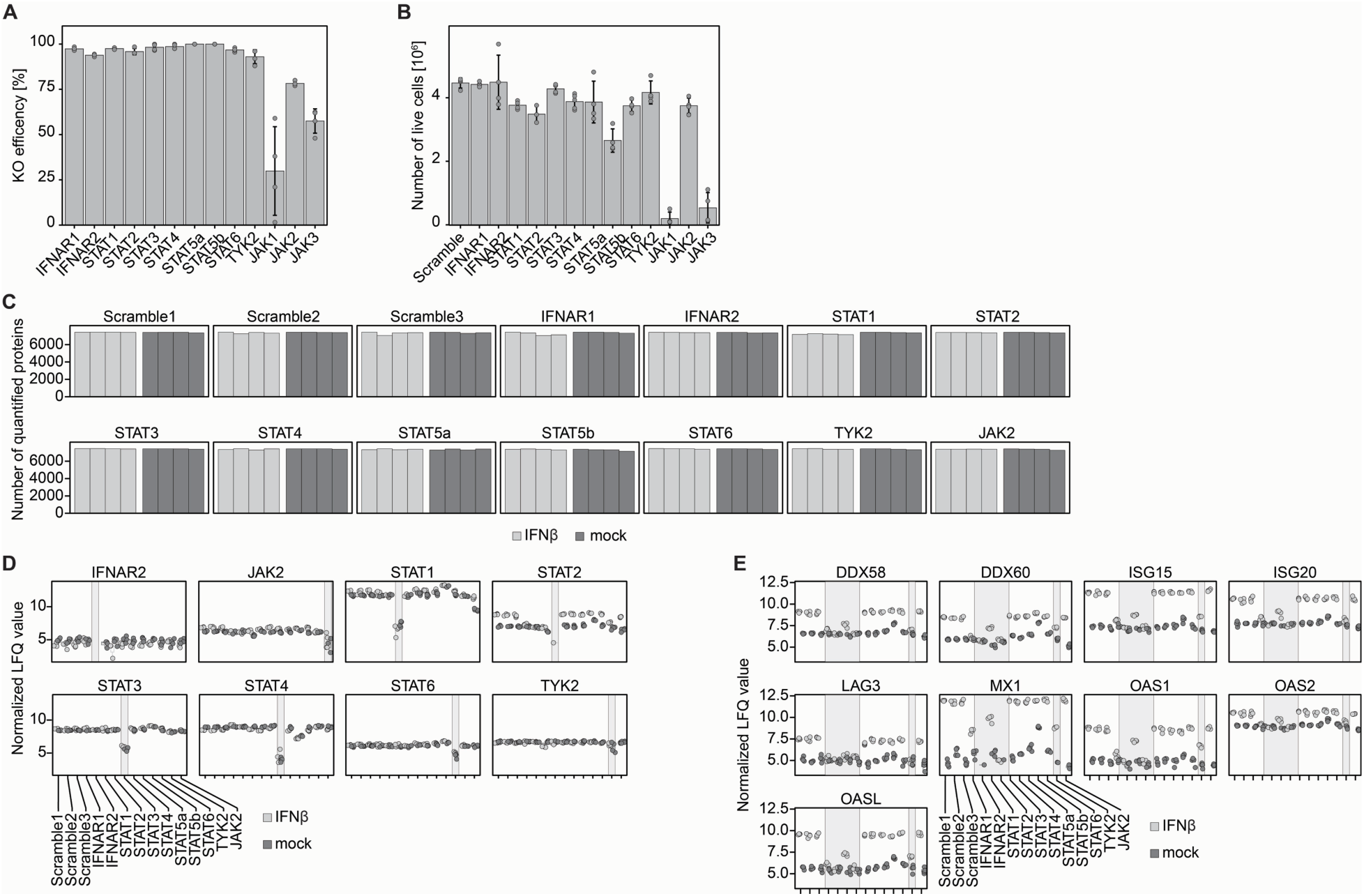
CRISPR-mediated gene knockout in primary CD4 T cells coupled with global proteomics identifies key regulators of the type-I interferon response. **(A)** Editing efficiency of individual CRISPR knockouts in primary CD4 T cells. (**B**) Number of viable cells per million after genetic editing. **(C)** Number of quantified proteins detected in each knockout sample. **(D)** Boxplots showing normalized LFQ intensities of knockout target proteins across all knockout conditions. Conditions in which the target protein was disrupted are indicated with a grey background. **(E)** Boxplots of normalized LFQ intensities for selected canonical ISGs across knockout conditions. Knockouts with significant changes relative to scrambled controls are highlighted with a grey background.

**Supplementary Figure 5:**
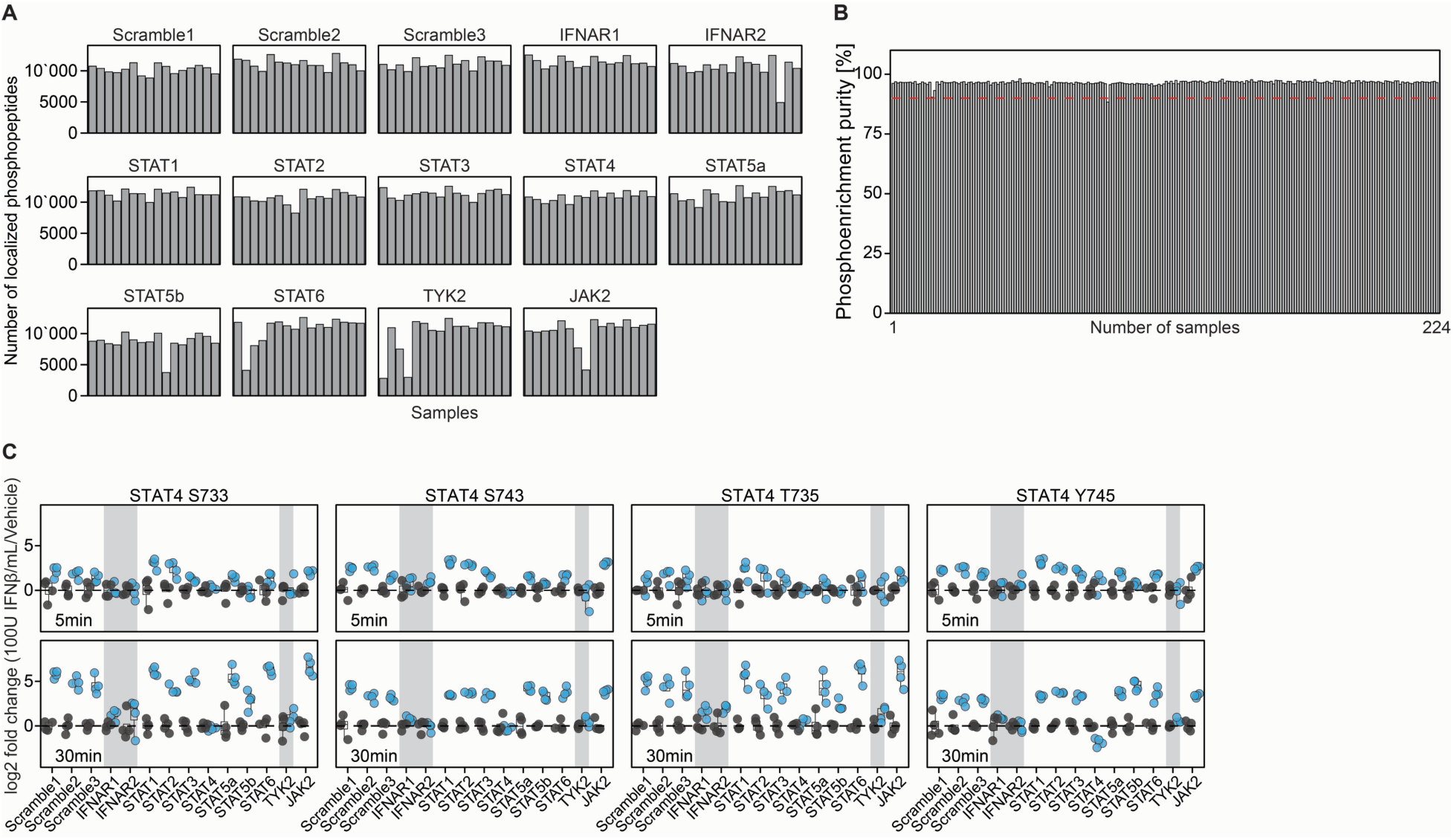
*simplePhos* enables scalable phosphoproteomic profiling in saturating CRISPR knockout primary CD4 T cells. (**A**) Number of localized phosphopeptides in each proteomics sample. (**B**) Purity of phosphopeptides across all samples; 90% threshold of phosphopeptide enrichment purity is shown in red. (**C**) Quantitative analysis of phosphorylation sites on STAT4 in CRISPR knockout CD4 T cells following interferon-β stimulation for 5 and 30 minutes. Phosphorylation levels are shown relative to vehicle-treated controls with matching genetic backgrounds. Knockouts with significant changes relative to scrambled controls are highlighted with a grey background.

**Supplementary Figure 6:**
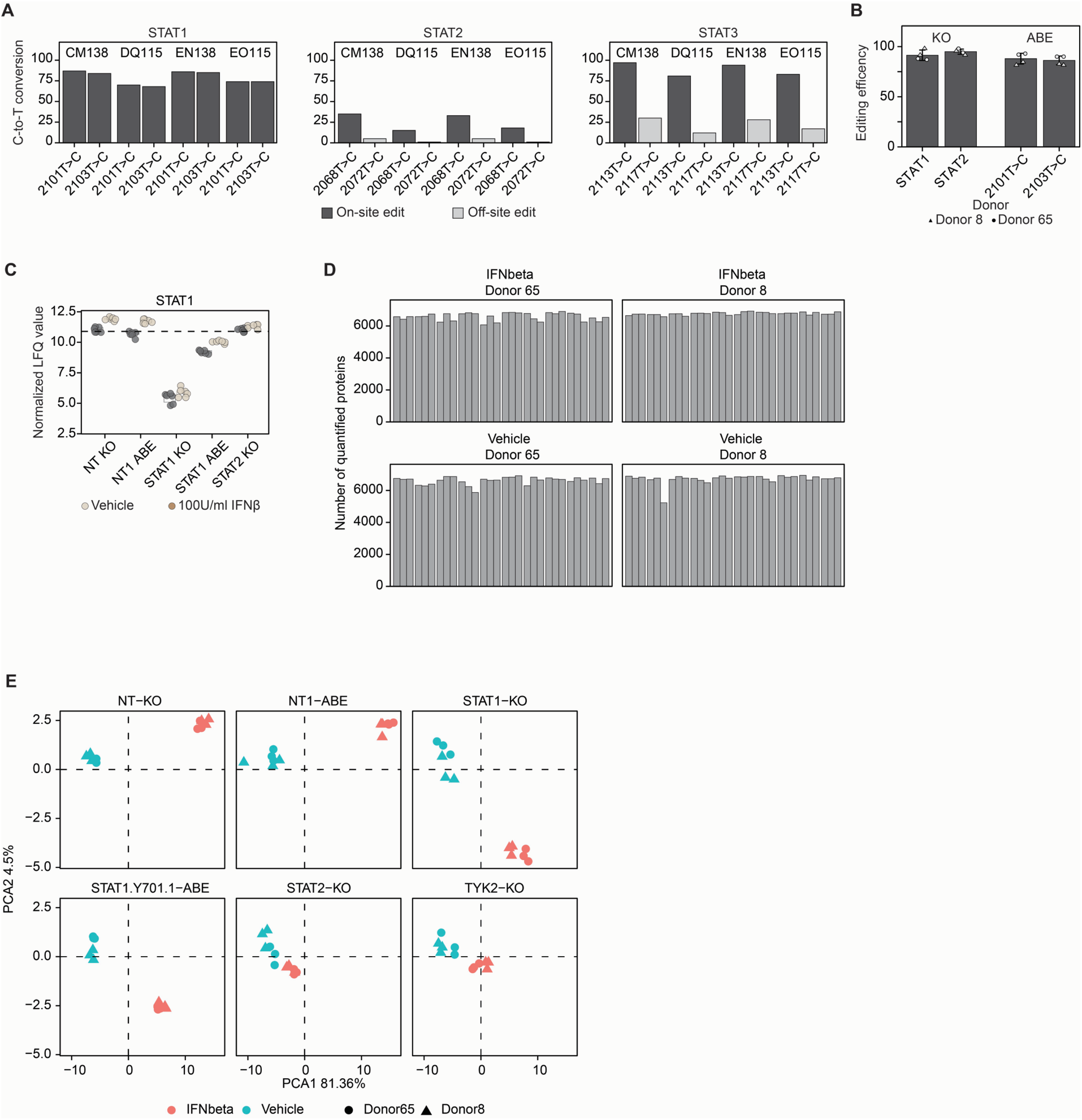
Base editing in primary CD4 T cells enables precise interrogation of signaling events. (**A**) C-to-T conversion rates at different loci within STAT1, STAT2, and STAT3 under various electroporation programs (Lonza 4D Nucleofector). On-target edits are shown in dark, and off-target edits are shown in grey. (**B**) Editing efficiency of STAT1 and STAT2 knockout samples, as well as STAT1 Y701H base-edited samples, across two independent donors. (**C**) LFQ of STAT1 protein abundance across different genetic perturbations, knockout or base editing. (**D**) Number of quantified proteins in each proteomics sample. (**E**) Principal component analysis (PCA) based on significantly regulated proteins following IFN-β stimulation in the various genetically edited samples.

